# Computational model for human 3D shape perception from a single specular image

**DOI:** 10.1101/383174

**Authors:** Takeaki Shimokawa, Akiko Nishio, Masa-aki Sato, Mitsuo Kawato, Hidehiko Komatsu

## Abstract

In natural conditions the human visual system can estimate the 3D shape of specular objects even from a single image. Although previous studies suggested that the orientation field plays a key role for 3D shape perception from specular reflections, its computational plausibility and possible mechanisms have not been investigated. In this study, to complement the orientation field information, we first add prior knowledge that objects are illuminated from above and utilize the vertical polarity of the intensity gradient. Then we construct an algorithm that incorporates these two image cues to estimate 3D shapes from a single specular image. We evaluated the algorithm with glossy and mirrored surfaces and found that 3D shapes can be recovered with a high correlation coefficient of around 0.8 with true surface shapes. Moreover, under a specific condition, the algorithm’s errors resembled those made by human observers. These findings show that the combination of the orientation field and the vertical polarity of the intensity gradient is computationally sufficient and probably reproduces essential representations used in human shape perception from specular reflections.

## Introduction

Specular reflections, which are seen in many daily objects, provide information about their material and surface finish [1-3], enhance the reality of animation and computer graphics, support 3D shape perception [4-6], and increase the 3D appearance of images [7].A specular reflection component in a single image can be regarded as a marking that is pasted on an object’s surface. However, the human visual system solves inverse optics, and we intuitively recognize that an image pattern is generated by a specular reflection [8].The regularity of the image patterns of specular reflections is closely related to 3D shape, and the human visual system perceives and evaluates specular reflection through coupled computation with 3D shape perception [9-11].

A previous psychophysical study showed that humans could recover 3D shapes from a single mirrored surface image under unknown natural illumination [12]. Furthermore, they hypothesized that the human visual system uses the orientation field for 3D shape perception from specular reflection and texture [12-14]. The orientation field is a collection of dominant orientations at every image location (Fig 1A), and this information is represented in the primary visual cortex (V1), which contains cells tuned to specific orientations [15]. In support of their hypothesis, they showed that 3D shape perception is modulated by psychophysical adaptation to specific orientation fields [13]. However, how 3D shapes are reconstructed from the orientation field, and whether it is adequate for 3D shape recovery remains unknown. Tappen [16] proposed a shape recovery algorithm and recovered the 3D shape of simple mirrored surfaces with curvature constraints by an orientation field from a single image under an unknown natural illumination. This suggests a possible mechanism of 3D shape perception from specular reflections. However, since the method is limited to convex shapes, it only explains a small part of human shape perception, which can recover more general shapes including both convex and concave regions [12].

**Fig 1.**
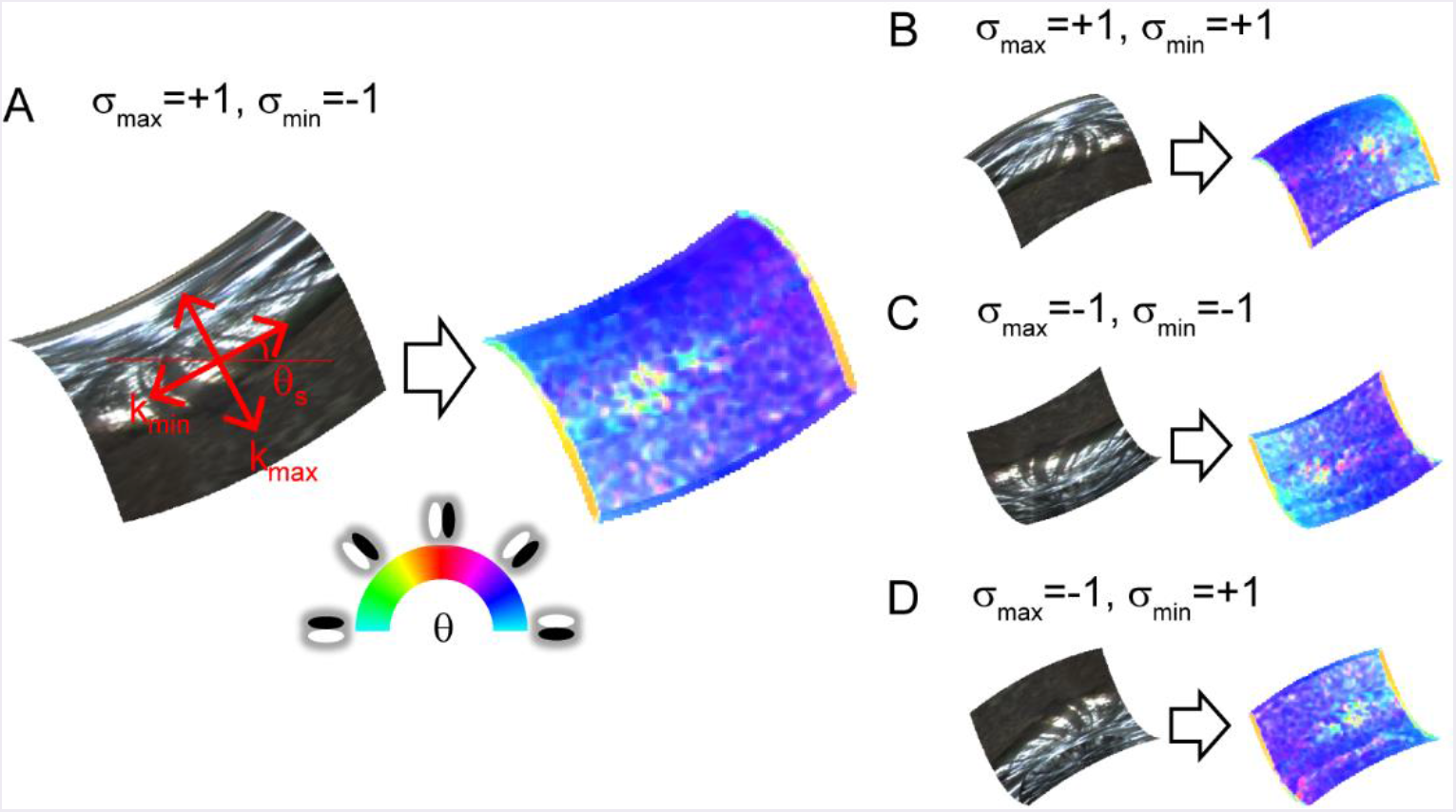
Orientation field of mirrored surface patches. Orientation fields are depicted on right side of images. Hue represents image orientation to which V1-cell-like oriented filter maximally responds at each location. Saturation represents degree of clarity of the image orientation (i.e., image anisotropy). (A) Surface second derivatives’ orientations of surface patch are explained in red on the mirrored surface. kmax and kmm represent large and small surface second derivative. θs represents surface orientation. σmax and σmm represent signs of kmax and kmm. (B), (C), (D) Surface patches have identical magnitude and orientation of surface second derivatives as A, but second derivative signs are different.

Other algorithms have also been proposed to recover 3D shapes from specular images. They employed either a known calibrated scene [17-19] or multiple images such as specular flows [20], motions of reflection correspondences [21], or line tracking [22]. Although they are useful in some situations, they cannot recover 3D shapes from a single specular image in an unknown scene. Li et al. recovered shapes using reflection correspondences extracted by SIFT [23] just using a single image under an unknown illumination environment like our proposed algorithm. However, their method is limited because it requires the known surface normal values of several surface points to constrain their results.

In this study, we recover general shapes containing both convex and concave surface regions using the orientation field. However, an innate problem prevents the recovery of general shapes from it. Here, we briefly explain the information of 3D shapes contained in the orientation field and its limitation as well as a strategy to overcome that limitation.

Fig 1 shows the relationship between the orientation field and the second order derivatives of the surface depth, which can be decomposed into two orthogonal orientations (left side of Fig 1A). This decomposed second derivatives are closely related to the principal curvatures, but these are not strictly the same (see Methods). The right side of Fig 1A represents the orientation field. In specular reflection, the illumination environment is reflected and appears in the image. At that time, the illumination environment is compressed toward a strong surface second derivative orientation and elongated along a weak surface second derivative orientation [12, 24]. As a result, image orientation θ is generated along small surface second derivative orientation θs. Moreover, the image anisotropy (the degree of the image orientation’s clarity, see Methods) also approximates the surface anisotropy (the ratio of the large and small surface second derivatives, see Methods) [12, 24]. The proposed algorithm uses this relationship for 3D surface recovery. Here, the problem is that the shape is ambiguous whether concave or convex, as shown in Fig 1B, 1C, and 1D. The image orientations are identical as Fig 1A because the surface orientations are also the same. However, the two signs of the surface second derivatives are different. The orientation field cannot distinguish among these four types.

We overcome the problem of concave/convex ambiguity by imposing a prior that illumination is from above [25, 26] (hereafter called the “above illumination prior”). In utilizing this prior knowledge, we actively use both a diffuse and a specular reflection component. Since most objects that give specular reflection also give diffuse reflection, a natural extension is to combine the features of both reflection components. Note that this prior also works for mirrored surfaces (see the Results section) and the human performance to resolve the concave/convex ambiguity from a mirrored surface increased when the illumination environment was brighter in the upper hemisphere [27].

We propose using the vertical polarity of the intensity gradient (hereafter vertical polarity) as an image cue (Fig 2; see also Supplementary Note1 in S1 Text). As with the orientation field, vertical polarity can be obtained by a V1-like filter [28] and its relation with 3D shape perception was reported [29]. Assuming the above illumination and Lambert reflectance, vertical polarity corresponds to the surface second derivative sign of vertical orientation (see Method). This prior is used only as an initial value for the optimization for 3D shape recovery. Because physically possible shape patterns given the orientation field are restricted [30, 31], it is expected that the remaining ambiguity (i.e., the surface second derivative sign of the horizontal orientation) is implicitly resolved and erroneous initial values are corrected through optimization.

**Fig 2.**
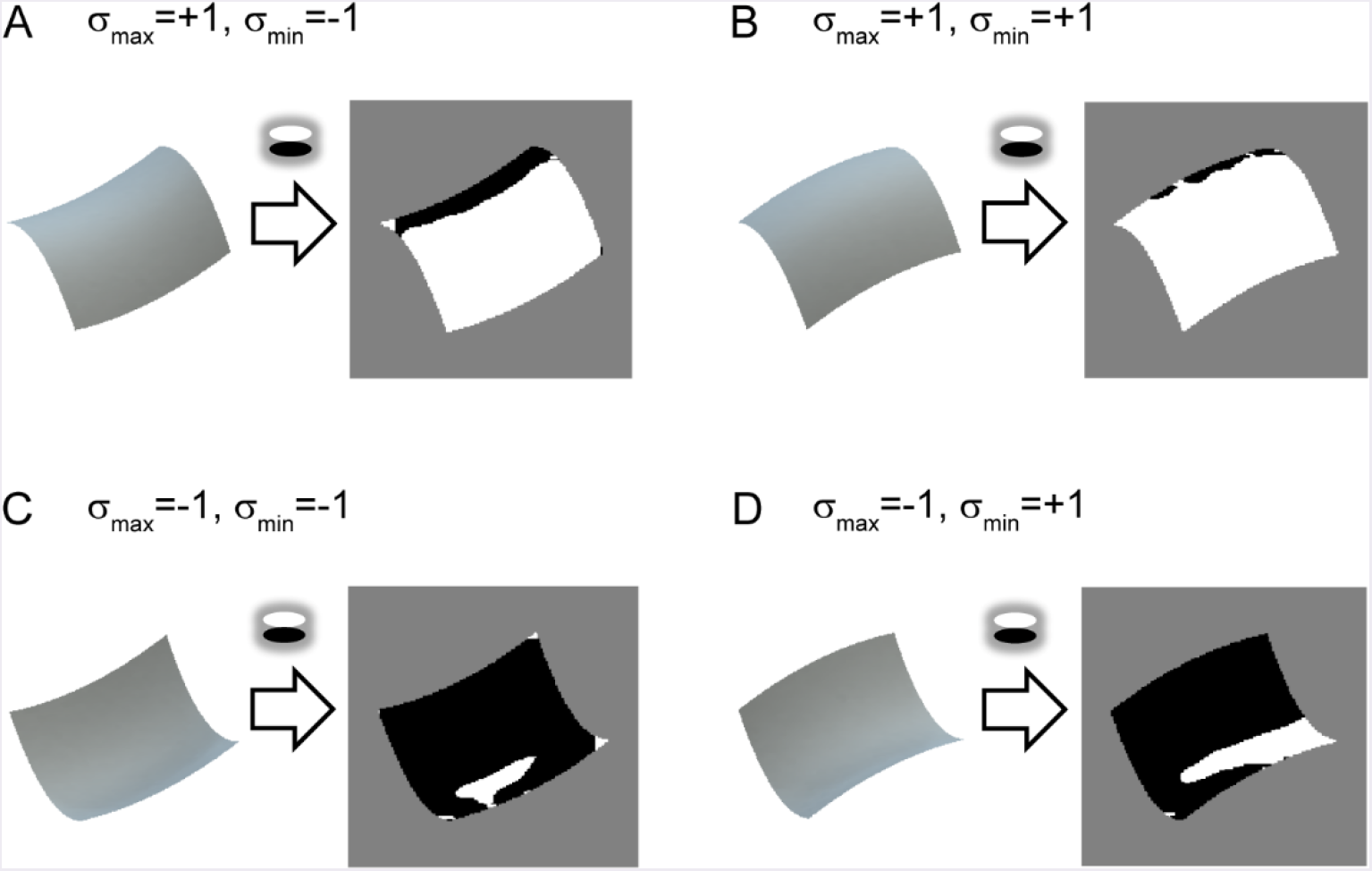
Relationship between vertical polarity and surface second derivative sign. Shaded images of identical surface patches to Fig. 1 are shown on left. Vertical polarity of each shaded image, obtained by extracting a sign of oriented filter response of vertical direction, is depicted on right. White represents positive and black represents negative.

Our proposed algorithm, which can recover general shapes including both convex and concave regions under an unknown natural illumination, is based on the information used by the human visual system. Therefore, it makes a critical contribution to understanding the mechanism of 3D shape perception from specular reflections.

## Results

### Flowchart of proposed algorithm

Fig 3 shows the flowchart of the proposed algorithm that recovers the 3D surface depth from a single specular image. The main procedure is as follows. First, the orientation field is extracted from an image; second, the cost function is formulated based on the orientation field; finally, the 3D shape is recovered by minimizing the cost function. Additionally, we extracted the vertical polarity from the image to resolve the concave/convex ambiguity. The initial values of the surface second derivative signs, σmax and σmin, are calculated based on the vertical polarity and used to minimize the cost function. The boundary conditions are also used, although they are omitted from this flowchart. The boundary conditions to resolve the ambiguity about the translation and affine transformation are incorporated in the cost function. The curvature sign of the 2D contour is calculated to obtain the signs of the 3D surface second derivative near the boundary. The details are described in Methods. The proposed algorithm outputs not only the recovered 3D surface depth but also the estimated surface second derivative signs, σmax and σmm, due to minimizing the cost function.

**Fig 3.**
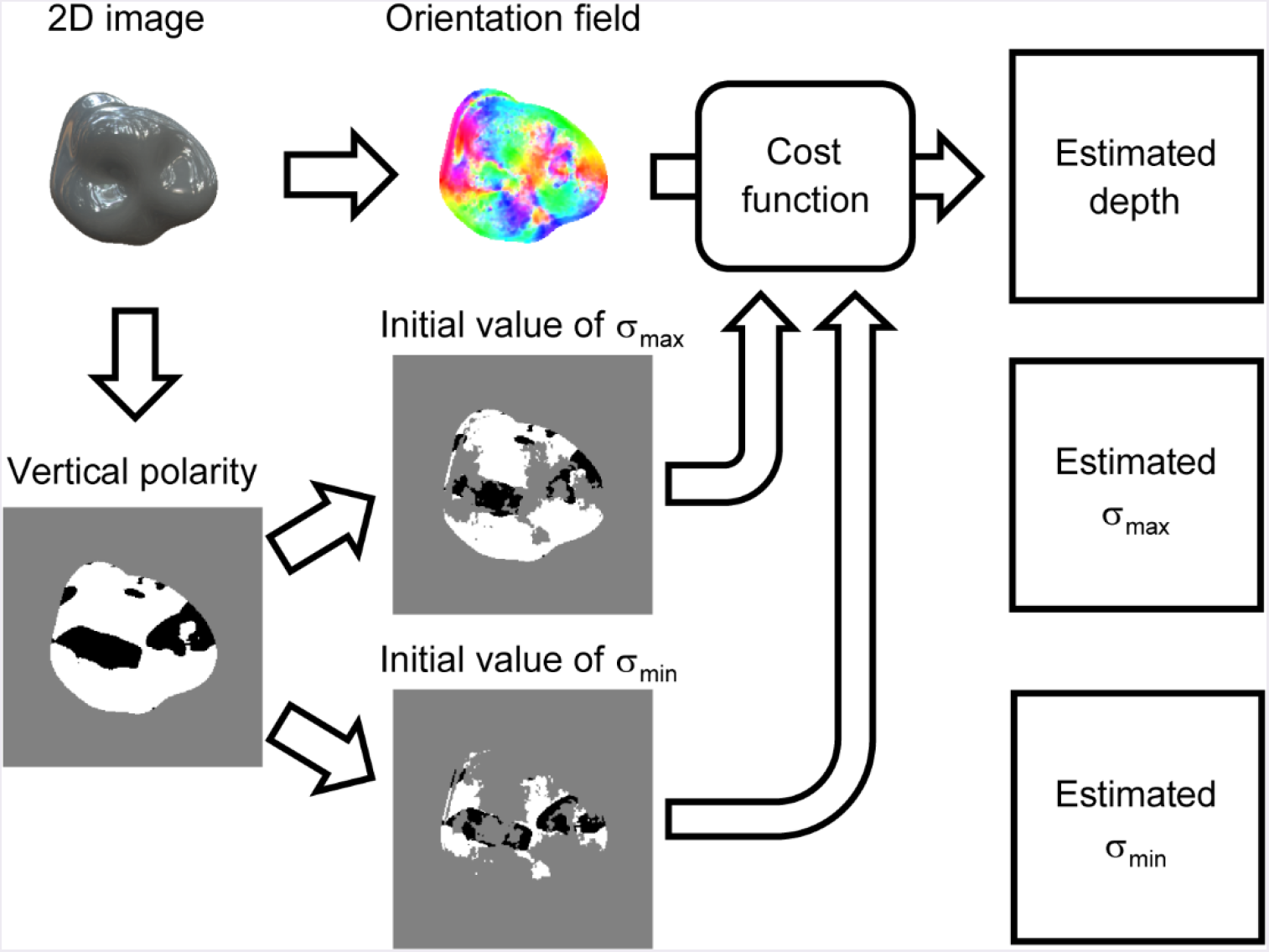
Flowchart of proposed shape recovery algorithm. Orientation field and vertical polarity are extracted from an image. Cost function is formulated based on orientation field. Initial values of signs of surface second derivative, σmax and σmm, are obtained by dividing vertical polarity. Estimated surface depth, σmax, and σmm are obtained by minimizing cost function.

### Shape recovery of glossy surfaces

The twelve glossy surfaces used to validate our proposed algorithm are shown in Fig 4. We generated them by computer graphics assuming both specular and diffuse reflections of the object’s surface (see Methods for details). The recovered shapes from these glossy surfaces are shown in Fig 5. The recovered depths are represented in grayscale; nearer surfaces are lighter and more distant surfaces are darker. Additionally,15 contour lines are superimposed. The estimated surface second derivative signs, σmax and σmm, are shown in S6 and S7 Figs. The true surface shapes and the true signs of the surface second derivative are shown in S1-S3 Figs.

**Fig 4.**
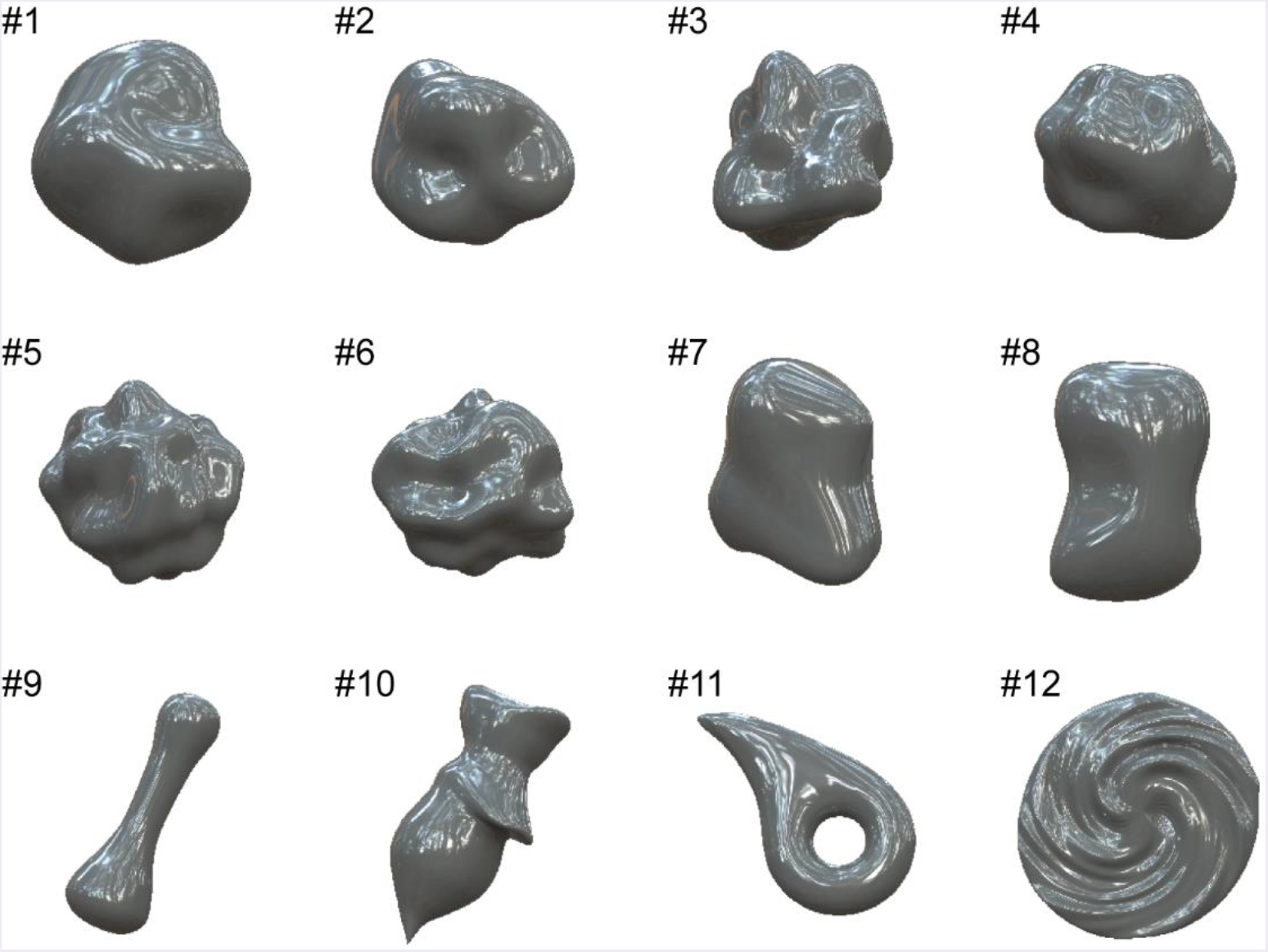
Glossy surfaces used to validate our proposed 3D shape recovery algorithm. These surfaces were generated by computer graphics assuming both specular and diffuse reflection on object’s surface.

**Fig 5.**
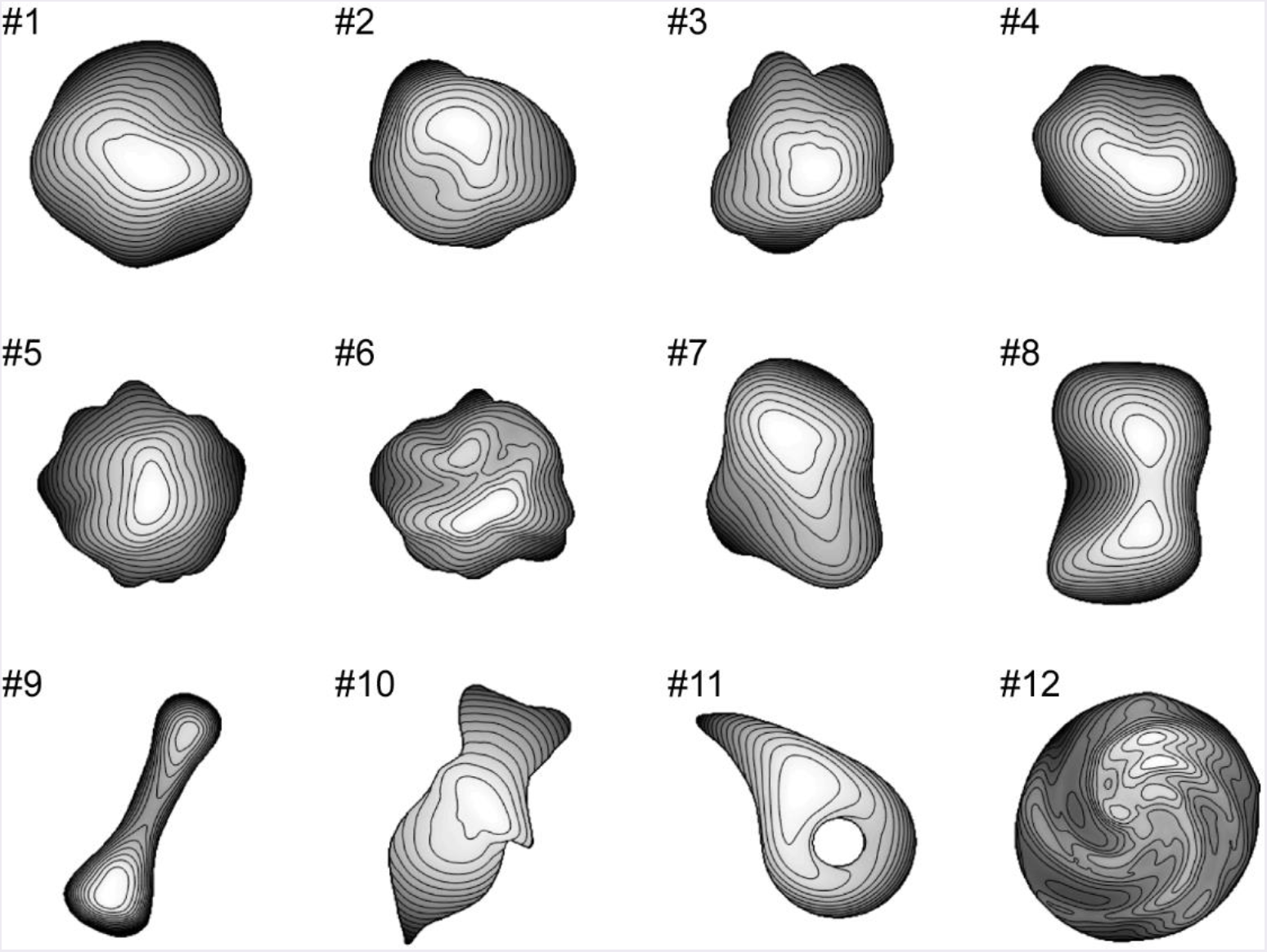
Recovered 3D shapes from glossy surfaces. Recovered surface shapes are represented by depth map and contour lines.

We evaluated the image cues (i.e., orientation field and vertical polarity) and the estimation results as follows. The orientation field error was quantified by the mean absolute errors throughout the object region between the image and surface orientations and between the image and surface anisotropies. We quantified the error of the vertical polarity by the correct ratio between the initial and true values of σmax and σmin, where the initial values exist. The shape recovery performance was quantified with two measures: global depth correlation rg and local interior depth correlation rn. The global depth correlation is simply the correlation coefficient of the recovered and true depths throughout the object region. The local interior depth correlation is the averaged value of the correlation coefficients of the recovered and true depths calculated in the local regions except near the boundary. The local interior depth correction is more sensitive to the agreement of the concavity and convexity inside the object region than the global depth correlation. Note that both depth correlations are calculated after the affine transformation so that the slant of the true surface depth becomes zero, because there is ambiguity about the recovered shape’s affine transformation [32]. No values were obtained of the local interior depth correlation of objects #9 and #11 because most of the object region is near the boundary. The details of the measures are described in Methods. The estimation performance of the surface second derivative signs, σmax and σmin, was quantified by the correct ratio with true values throughout the object region.

The average values of the mean absolute error of the orientation and anisotropy for 12 objects were 11.3^?^ and 0.15. The average values of the correct ratio of the initial values of σmax and σmin for 12 objects were 0.79 and 0.70. The initial values and the correct ratios of all objects are shown in S4 and S5 Figs.

The shape recovery performances of 12 objects (#1, #2,… #12) were as follows: global depth correlation rg = 0.98, 0.91, 0.87, 0.82, 0.89, 0.88, 0.90, 0.95, 0.89, 0.65, 0.65, 0.80 (average rg = 0.85); local interior depth correlation ru = 0.97, 0.71, 0.67, 0.66, 0.91, 0.86, 0.84, 0.95, -, 0.45, -, 0.60 (average ru = 0.76). As an impression of appearance, the shape recovery seems successful if both the global and local interior depth correlations exceed 0.7. The recovered shapes of objects #1, #2, #5, #6, #7, #8, and #9 resemble the 3D surface impressions received from the corresponding images in Fig 4. The global depth correlations of #10 and #11 and the local interior depth correlations of #3, #4, #10, and #12 were below 0.7. The recovered shapes of #3, #4, #11, and #12 were roughly good but lacked accuracy. The shape of object #10 was not well recovered. The following are the estimation performances of the surface second derivative signs: the correct ratios of the estimated σmax were 0.90, 0.90, 0.80, 0.81, 0.79, 0.86, 0.87, 0.93, 0.99, 0.86, 0.98, 0.70 (average 0.86); the correct ratios of the estimated σmin were 0.77, 0.74, 0.63, 0.68, 0.64, 0.68, 0.69, 0.82, 0.82, 0.75, 0.78, 0.60 (average 0.72). The correct ratios of the estimated σmax and σmin exceeded those of the initial values even though the initial values exist only in half of the object region.

### Shape recovery of mirrored surfaces

The proposed algorithm is applicable to mirrored surfaces without shading although we assumed that shading exists to obtain good initial values of the surface second derivative signs by calculating the vertical polarity. Fig 6 shows the mirrored surfaces used to validate our proposed algorithm.

**Fig 6.**
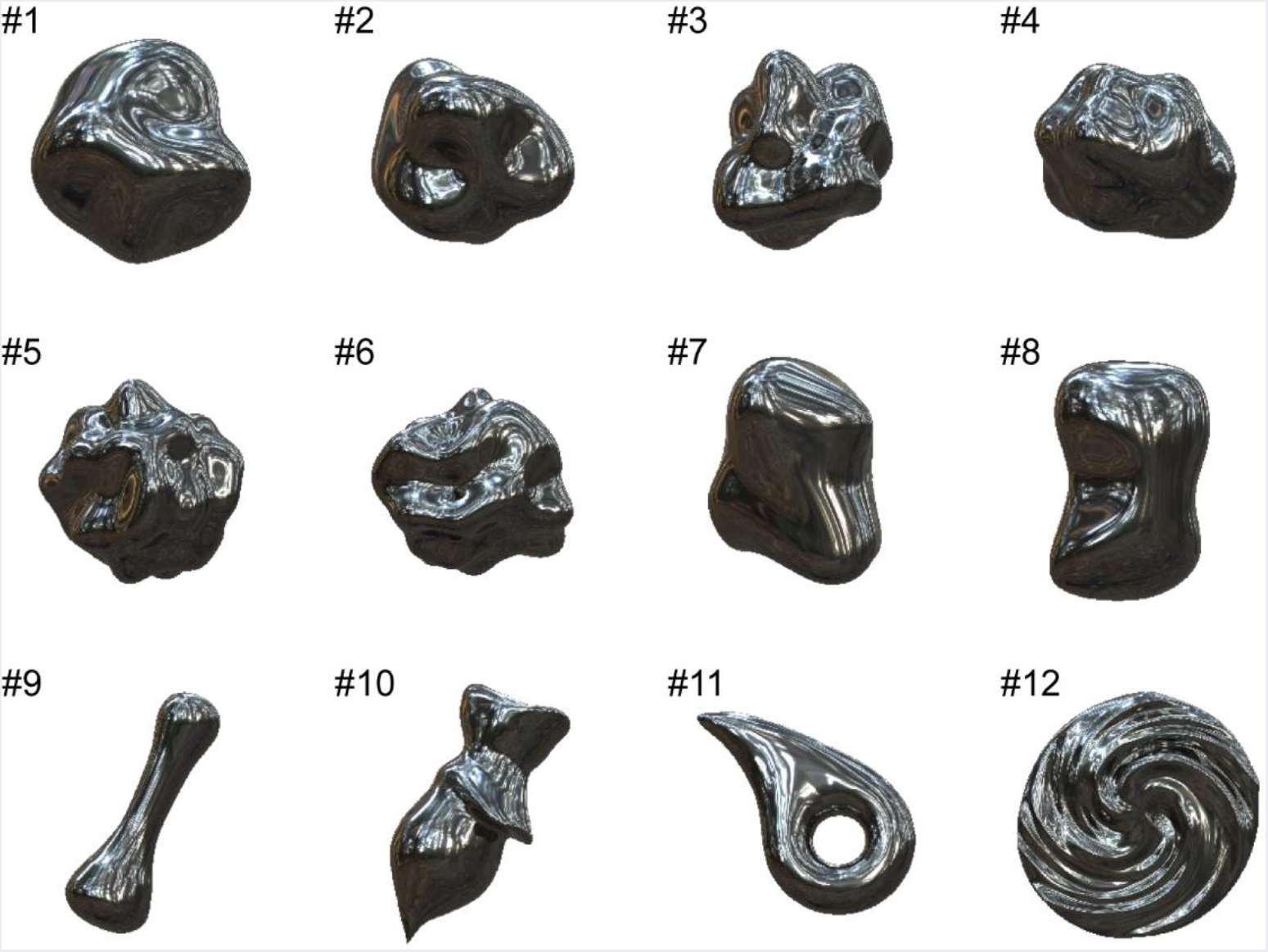
Mirrored surfaces used to validate our proposed 3D shape recovery algorithm. These surfaces were generated by computer graphics assuming only specular reflection on object’s surface.

The average values of the correct ratio of the initial values of σmax and σmm for 12 objects were 0.64 and 0.62. These correct ratios were significantly lower than those of the glossy surfaces, but still higher than a chance level of 0.5. The initial values and the correct ratios of all the objects are shown in S8 and S9 Figs. The average values of the mean absolute error of the orientation and the anisotropy for 12 objects were 10.9° and 0.13. These orientation field errors were slightly lower than those of the glossy surfaces, suggesting that the shading component slightly disturbed the relationship between the orientation field and the surface second derivative based on specular reflections.

The recovered shapes from the mirrored surfaces are shown in Fig 7. The following are the shape recovery performances: global depth correlation rg = 0.95, 0.93, 0.78, 0.81, 0.89, 0.89, 0.81, 0.93, 0.91, 0.66, 0.66, 0.86 (average rg = 0.84); local interior depth correlation rli = 0.96, 0.74, 0.60, 0.67, 0.90, 0.85, 0.67, 0.91, -, 0.55, -, 0.65 (average ru = 0.75). Although the appearances of the recovered shapes from the mirrored surfaces look noisier than those from the glossy surfaces (e.g., #1 and #8), the averaged global and local interior depth correlations differ by only 0.01 and 0.01, indicating that the proposed shape recovery algorithm is applicable to both mirrored and glossy surfaces. The following are the estimation performances of the surface second derivative signs. The correct ratios of the estimated σmax were 0.74, 0.85, 0.74, 0.72, 0.74, 0.79, 0.80, 0.84, 0.97, 0.77, 0.99, 0.68 (average 0.80); the correct ratios of the estimated σmm were 0.67,0.73, 0.64, 0.68, 0.63, 0.67, 0.68, 0.79, 0.83, 0.70, 0.79, 0.61 (average 0.70). The noisier appearance of the recovered shapes of the mirrored surfaces is related to the lower correct ratio of the estimated σmax than that of the glossy surfaces. The estimated surface second derivative signs, σmax and σmm, are shown in S10 and S11 Figs.

**Fig 7.**
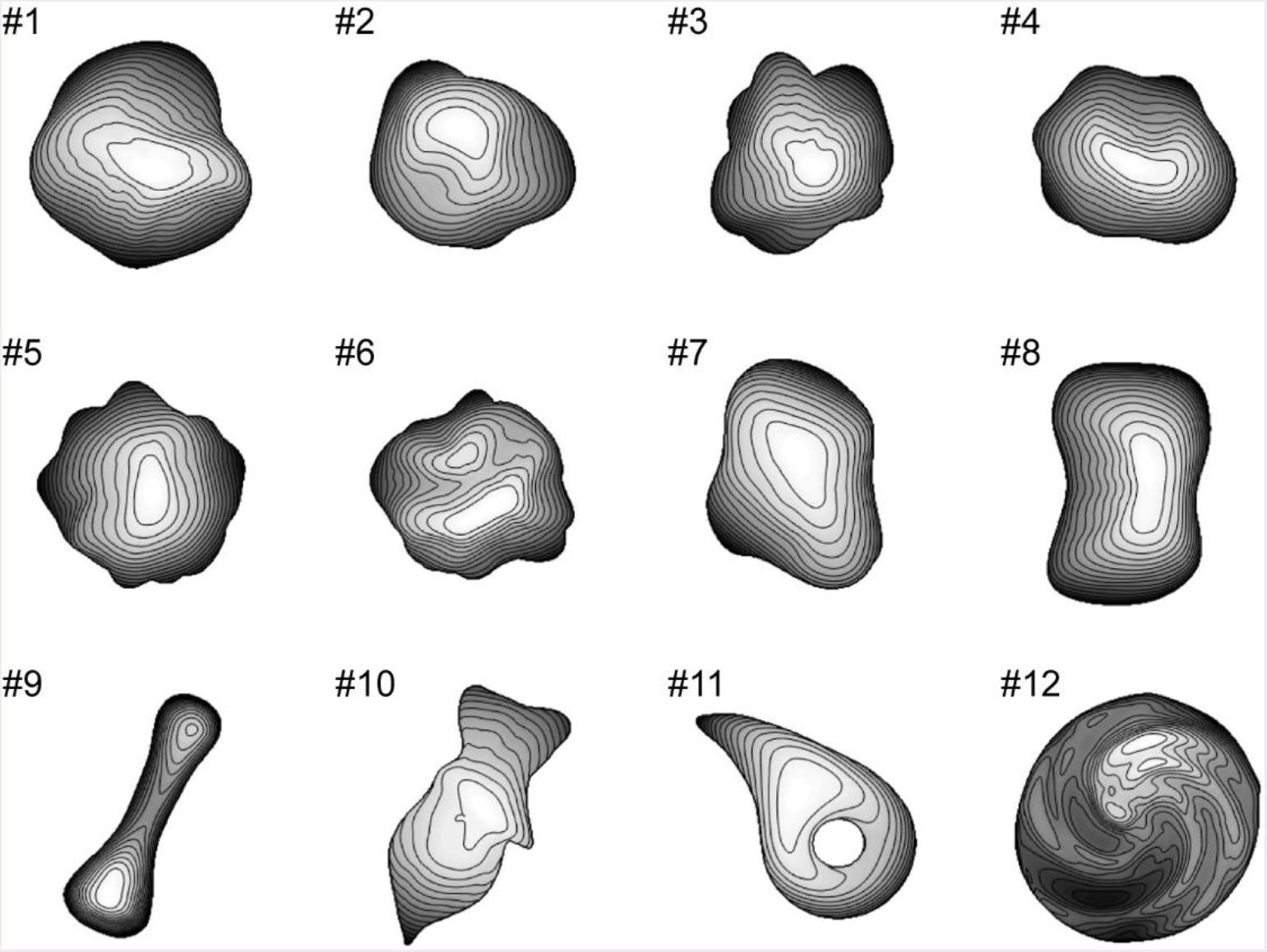
Recovered 3D shapes from mirrored surfaces. Recovered surface shapes are represented by depth map and contour lines.

### Estimation accuracy in different conditions

We tested the proposed algorithm in four different conditions. The first and second conditions are the shape recoveries from the glossy and mirrored surfaces shown in Figs 4 and 6 (denoted as glossy and mirrored conditions). In the third condition, the shapes were recovered from the glossy surfaces shown in Fig 4, but the above illumination prior was not used (denoted as the noAIP condition). And in the fourth, the shapes were recovered from the shape orientation fields that were obtained from the true 3D shapes (denoted as the shapeOF condition). Note that in the shapeOF condition, the same initial values of σmax and σmin were used as the glossy condition. Tables 1 and 2 summarize the errors of the image cues and the estimation performances of the four conditions. Additionally, we tested the algorithm in three more conditions to investigate the effect of the contour constraint, the illumination environment, and the image resolution. These results are shown in Supplementary Note 2 in S1 Text.

**Table 1.** Errors of image cues.

|  | Mean absolute error |  | Correct ratio |  |
| --- | --- | --- | --- | --- |
| | Orientation | Anisotropy | Initial $\sigma_{\max}$ | Initial $\sigma_{\min}$ |
| Glossy | 11.3° | 0.15 | 0.79 | 0.70 |
| Mirrored | 10.9° | 0.13 | 0.64 | 0.62 |
| noAIP | (11.3°) | (0.15) | 0.85 | 0.67 |
| shapeOF | 0° | 0 | (0.79) | (0.70) |

Orientation field errors and correct ratios of initial values of $\sigma_{\max}$ and $\sigma_{\min}$ , which are
averaged values of 12 objects. Values of orientation field errors of noAIP condition and correct ratios of initial values of shapeOF condition are parenthesized because these are identical as glossy condition.

**Table 2.** Estimation performances.

|  | Shape recovery accuracy |  | Correct ratio |  |
| --- | --- | --- | --- | --- |
| | $r_g$ | $r_{li}$ | Estimated $\sigma_{\max}$ | Estimated $\sigma_{\min}$ |
| Glossy | 0.85 | 0.76 | 0.86 | 0.72 |
| Mirrored | 0.84 | 0.75 | 0.80 | 0.70 |
| noAIP | 0.77 | 0.52 | 0.80 | 0.67 |
| shapeOF | 0.87 | 0.88 | 0.92 | 0.80 |

Global and local interior depth correlations of recovered shapes and correct ratios of estimated signs of surface second derivative. These are averaged values of 12 objects.

In the noAIP condition, the shapes were recovered from the glossy surfaces without the above illumination prior to check its necessity. In this condition, the initial values of σmax and σmm were all set to +1 based on the convex prior possessed by humans [33, 34]. The average values of the correct ratio of the initial values of σmax and σmm for 12 objects were 0.85 and 0.67. First, the shapes were recovered with the same algorithm that was used with the other conditions. As a result, the estimated σmax and σmm were almost the same as the initial values; 98% and 88% of the estimated σmax and σmm were +1. This means that the estimation failed. The average values of the global and local interior depth correlations for 12 objects were rg = 0.74 and rn = 0.48. These estimation performances are not summarized in Table 2, because the estimation completely failed. Next we altered the temperature parameter of the mean field algorithm (see Supplementary Note 3 in S1 Text for details) from ß0=10 to ß0=1 to extend the search range, since the initial values were not reliable in this condition. As a result, we obtained better shape recovery results. The average values of the global and local interior depth correlation for 12 objects were rg = 0.77 and rli = 0.52. The average values of the correct ratio of the estimated σmax and σmm for 12 objects were 0.80 and 0.67. The estimation performances of objects #1, #8, and #9 were high despite the noAIP condition. However, most of the recovered shapes look noisy, probably because of the alternation of the temperature parameter, and the estimation performance was lowest in the four conditions. The recovered shapes and the estimated signs of the surface second derivative of the noAIP condition are shown in S12-S14 Figs.

In the shapeOF condition, the shapes were recovered from the surface orientations that were obtained from the true 3D shapes instead of the image orientations to investigate the effect of the orientation field errors on the shape recovery errors. In this condition, the vertical polarity of the glossy surfaces was used to resolve the concave/convex ambiguity. The average values of the global and local interior depth correlations for 12 objects were rg = 0.87 and rli = 0.88. The average values of the correct ratio of the estimated σmax and σmm for 12 objects were 0.92 and 0.80. The estimation performances of the shapeOF condition were very high, except for objects #9 and #10, and significantly higher than the other conditions. The recovered shapes and the estimated signs of the surface second derivative of the shapeOF condition are shown in S15-S17 Figs.

### Consistency with human shape perception

Finally, we conducted a psychophysical experiment to investigate the linkage between the shape recovery algorithm and human shape perception. We prepared a glossy surface image that evokes 3D shape misperception (Fig 8A) by using another illumination environment that is inconsistent with the above illumination prior (Galileo’s Tomb of the Devebec dataset) and carefully modifying the 3D object’s shape. Fig 8B is an image of the same object rendered under identical illumination environments as Figs 4 and 6 (Eucalyptus Grove of the Devebec dataset). Fig 8C represents the depth map of the true 3D shapes. The red cross indicates where the surface looks concave from Fig 8A, although the surface looks convex from Fig 8B and the true surface is convex. Fig 8D and 8E indicate the recovered shapes from the images of Fig 8A and 8B. In accordance with the appearance, the recovered shape from Fig 8A is concave and that from Fig 8B is convex around the red cross mark. The estimation performances (rg, r¾ correct ratio of estimated σmax and correct ratio of estimated σmin) of Fig 8D and 8E were (0.91, 0.76, 0.73, 0.60) and (0.98, 0.99, 0.90, 0.84).

**Fig 8.**
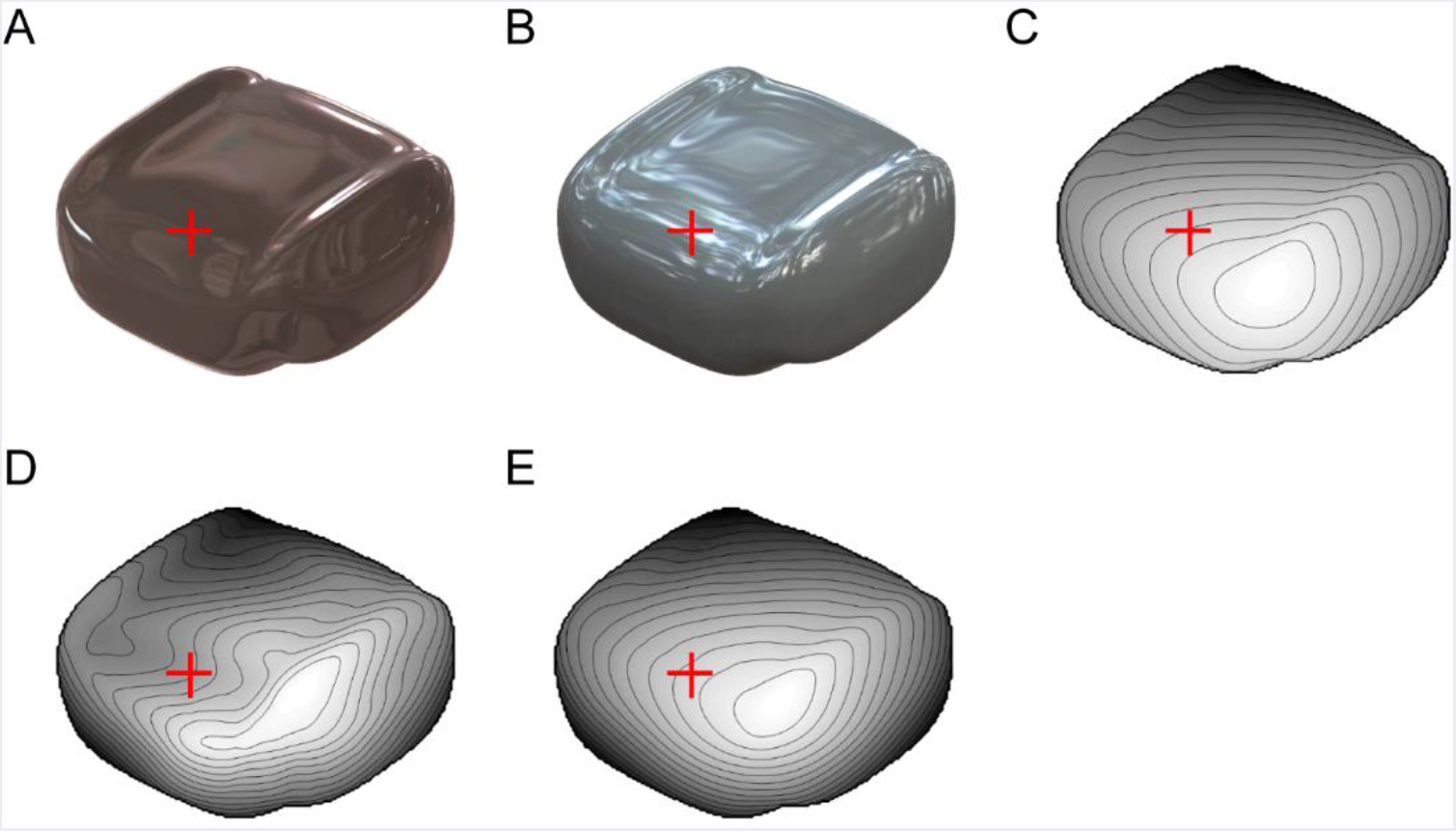
Images used for psychophysical experiment. (A) Glossy surface rendered in indoor environment. Red crosses indicate position where misperception likely occurs. (B) Glossy surface of identical object as A rendered in outdoor environment. (C) Depth map of true 3D shapes of *A* and *B*. (D) Recovered shape from image in A. (E) Recovered shape from image in *B*.

In psychophysical experiments, five subjects were first asked whether the local 3D surface around the red crosses in Fig 8A and 8B looks convex or concave. After that, they were asked whether the true 3D shape (Fig 8C), the recovered 3D shape (Fig 8D, or 8E) was more similar to the perceived 3D shape from the image. Four of five subjects answered that the local surface of Fig 8A looked concave and only one thought that it looked convex. All five subjects answered that the local surface of Fig 8B looked convex. Four of five subjects answered that the recovered shape (Fig 8D) was closer to the perceived shape of the image shown in Fig 8A, and one thought that the true shape (Fig 8C) was closer. Four of five subjects answered that the recovered shape (Fig 8E) was closer to the perceived shape from the image shown in Fig 8B and one answered that the true shape was closer. To summarize, most subjects (4 of 5) perceived the incorrect shape from Fig 8A and the recovered shape (Fig 8D) was consistent with the misperceived shape

## Discussion

We developed an algorithm that estimates 3D shapes from a single specular image to investigate a possible mechanism of human 3D shape perception from specular reflections. This algorithm mainly relies on the orientation field suggested by a previous psychophysical study [12]. However, since the orientation field cannot resolve the local concave/convex ambiguity, the 3D shape recovery from it alone was difficult (see the noAIP condition, Table 2). To resolve the concave/convex ambiguity, we added the prior knowledge that objects are illuminated from above. The vertical polarity of the intensity gradient is an image cue to utilize this prior knowledge. We evaluated the developed algorithm with the glossy and mirrored surfaces of 12 complex shapes. The depth correlations between the recovered and the true shapes were as high as around 0.8. To further confirm the necessity of the vertical polarity information, we also conducted a psychophysical experiment with an image that caused human misperception due to the inconsistency with the above illumination prior. The human-misperceived and recovered shapes were consistent in most subjects. These findings show that the vertical polarity of the intensity gradient as well as the orientation field are related to 3D shape perception and the combination of both enables 3D shape recovery from a single specular image.

### Shape recovery of mirrored surfaces

The shape recovery performance of the mirrored condition was almost as high as the glossy condition (Table 2), although the relationship between the vertical polarity and the surface second derivative sign was only proved in the diffuse reflection component (see Methods). The present result indicates that vertical polarity of the specular component was also useful for the initial second derivative signs for the following reason. The diffuse reflectance component in Fig 2 shows a relationship where the luminance is high in the upper side and low in the lower side when the surface is convex with respect to the vertical orientation (Fig 2A and 2B) and vice versa (Fig 2C and 2D). The same relationship holds for the mirrored surfaces of Fig 1. The luminance tends to be higher in the upper side than the lower side when the surface is convex (Fig 1A and 1B) and vice versa (Fig 1C and 1D). Thus, the vertical polarity of the mirrored surface at low frequencies is related to the surface second derivative sign of the vertical orientation, although the high-frequency component is not related to it. When the vertical polarity is calculated, a relatively low-frequency image component is extracted and further smoothed to remove the high-frequency component of the specular reflection (see Methods). Therefore, it provides meaningful information about second derivative signs even from mirrored surfaces, although the correct ratio of the initial sign values of the mirrored condition is actually worse than that of the glossy condition (Table 1).

### Representation of surface curvatures

In this study, the sign and magnitude of the surface second derivatives are separately described. Similar representation can be seen in some psychophysical experiments [35, 36], in which subjects classified 3D shapes based on curvature signs. Furthermore, the neural representation of surface curvatures was studied in electrophysiological experiments. Srivastava et al. showed that the neurons in the inferior temporal cortex (the area for object recognition) of macaques are mainly sensitive to the curvature sign,but the neurons in the anterior intraparietal area (the area for motor planning) are sensitive to the curvature magnitude as well as the sign [37]. This might suggest that the curvature sign’s representation is important for object recognition, and its magnitude is also required for motor planning. These and other psychophysical and electrophysiological studies [38, 39] provide hints to develop more efficient and human-like shape recovery algorithms.

The estimation of a small surface second derivative sign, σmin, was more difficult than that of a large surface second derivative, σmax, in all four conditions (see right half of Table 2). A similar phenomenon can be seen in human shape perception. When subjects classified local shapes based on the curvature signs, saddles were often misclassified as ridges or ruts (convex or concave cylinders) [35, 36], suggesting that humans often neglect the small surface curvature of saddle shapes. Since the small surface curvature is less visible in the image, its estimation is intrinsically difficult. In the proposed algorithm, the small second derivative sign is forcibly classified as +1 or −1, but it might be better to treat it ambiguously like the quantum superposition when its classification is difficult.

Note here that the shape recovery from specular reflections has much in common with that from line drawings because lines or specular orientations appear at the high curvature in both cases [40, 41]. In a line drawing study, edge-labeling algorithms classified the orientation edges as either convex or concave [30, 31]. This corresponds to the determination of the large surface second derivative sign in our study. It would be interesting to find and utilize the similarities of the shape recovery algorithms from specular reflection and line drawing [42].

### Origin of shape recovery errors

The orientation field error is a major error factor of the proposed algorithm, because the shape recovery performance was very high in the shapeOF condition (Table 2 and S15 Fig). In this condition, the surface second derivative signs were accurately estimated even though the initial values from the vertical polarity were somewhat incorrect and absent in half of the region. This result indicates that the proposed shape recovery algorithm works well at least under such ideal conditions. Therefore, the error due to the proposed algorithm’s methodological imperfection is relatively small. It also indicates that the orientation field is satisfactory for the 3D shape recovery of such curved surfaces examined in this study with the help of the above illumination prior. The difference of the shape recovery performances between the glossy and shapeOF conditions reflects errors that originate from the image orientation field. Compared with the orientation field error, the effects of the initial second derivative sign errors are limited because they are expected to be corrected through optimization; orientation field error inevitably affects the resultant shape because it is directly incorporated in the cost function. Actually, the shape recovery performance of the mirrored condition was comparable to the glossy condition even though the initial second derivative sign errors of the mirrored condition were significantly larger than those of the glossy condition. Of course, too many initial errors cannot be corrected as suggested by the poor shape recovery performance of the noAIP condition. The orientation field errors probably affect the error corrections of the initial values through optimization.

### Interpretation of human shape misperception

The glossy surface Fig 8A, which was used for our psychophysical experiment, looks concave around the red cross mark, but Fig 8B looks convex. The illumination environment caused this difference. The Eucalyptus Grove environment for Fig 8B is outdoors and consistent with the above illumination prior of humans. However, the Galileo’s Tomb environment for Fig 8A is indoors and the ceiling is dark that is against the above illumination prior. The dark ceiling caused a negative value of the vertical polarity around the red cross mark despite its convex 3D shape, which presumably caused the concave interpretation. In this example, both the convex and concave interpretations are consistent with the surrounding information. Therefore, humans may interpret images as convex or concave depending on the illumination environment.

### Limitations and future work

The following are the limitations of our shape recovery algorithm. First, since it is based on the relationship between the orientation field and the surface second derivative, large error occurs when this relationship is invalid. For example, if the illumination environment is biased to a specific orientation (e.g., striped illumination), it biases the image orientation [24]. The orientation error becomes large where the surface anisotropy is small [24]. For example, if the true shape is a plane (i.e., the surface anisotropy is zero), the image orientation reflects not the surface second derivative but only the orientation of the illumination environment and causes shape recovery errors. Second, images under an unnatural illumination environment against the above illumination prior could not be properly recovered as it is difficult for humans [27, 43]. Third, the proposed algorithm cannot estimate the depth scale as well as the slant due to the ambiguity about the affine transformation of the recovered shape [32]. Humans also have difficulty estimating the slant [6, 44] and the depth scale [6, 32] from a single image without prior knowledge of the object’s shape. Therefore, we evaluated the recovered shapes by depth correlations after the affine transformation so that the slant of the true surface depth becomes zero. We did not evaluate the normal map because it depends on the depth scale. Fourth, because the proposed algorithm assumes that the surface depth is second order differentiable, it cannot properly treat bends, cusps, and self-occlusion inside the object region (occluding edges or limbs [31]) and generates smoother shapes than actual shapes. This property may worsen the shape recovery performance of objects #10, #3, and #4. Note that the limitations listed above (except for the fourth) are closely related to the limitations of human shape perception.

Future work has several promising directions. First, further psychophysical experiments are required to understand human shape perception from specular reflections in detail and will help improve the shape recovery algorithm to better simulate the human shape perception. It would be interesting to use the image-based shape manipulation method based on the orientation field [45] to compare the recovered and human-perceived shapes. Second, the proposed shape recovery algorithm will be useful for computer vision methods. By integrating it with a study that estimates material (BRDF) from a single image of a known shape [46], it might become possible to estimate an unknown shape’s material. By providing more accurate recovered depth information, we expect to enhance the reality of the image-based material editing that is based on shape information [47]. For further improvement of the shape recovery performance, the proposed shape from the specularity algorithm could be integrated with the shape from shading algorithms [48, 49], where it would be helpful to use color information to separate diffuse and specular reflection components [50]. Third, it would be interesting to study whether 3D shapes can be recovered from translucent images with specularities. A previous study [51] argued that an object looks translucent when images are manipulated so that the diffuse reflection component is contrast-reversed,but the specular reflection component is left intact. This result suggests that we must alter how the specular and diffuse reflection components are combined for shape recovery from translucent images, such as reversing the sign of the vertical polarity in the case of translucent images compared with opaque images.

## Methods

As a precondition to 3D shape recovery, we assume that the image region is known where the object exists. It may be obtained by an edge detection algorithm or decided by humans. We denote the object region as Ω, the number of pixels in Ω as N_Ω_, the boundary region, which is the region between the boundary of Ω and one pixel inside it, as ∂Ω, and the number of pixels in ðΩ as N∂Ω. The resolution of the 3D shape recovery was 256 × 256 pixels. We set a Cartesian coordinate on the image plane, where the x and y axes represent the horizontal and vertical axes of an image plane and the z axis represents the front direction. We represent the depth of the 3D object surface as z(x,y). The following notations are used: scalars are represented in normal-type letters as x; vectors are represented in lower-case boldface letters as **x**; matrices are represented in upper-case boldface letters as **X**.

### Images and extraction of image cues

We used the images of 12 different 3D shapes to evaluate the proposed algorithm (Figs 4 and 6). The images had 1024 × 1024 pixel resolution and were colored, although they were downsampled to 256 × 256 pixels before the 3D shape recovery and became achromatic because the proposed algorithm does not use color information. These images were rendered by Radiance software (http://radsite.lbl.gov/radiance/). The surface reflection property was modeled by the Ward-Duer model [52, 53]. We set diffuse reflectance pd, specular reflectance ρ_s_, and the spread of specular reflection α as pd = 0.1, p_s_ = 0.15, α = 0 for the glossy surfaces (Fig 4) and pd = 0, p_s_ = 0.25, α = 0 for the mirrored surfaces (Fig 6). For the natural illumination environment, we used a high dynamic range image from the Devebec dataset (http://ict.debevec.org/~debevec/; Eucalyptus Grove). For the quadratic patch images in Figs 1 and 2, we set pd = 0, p_s_ = 0.25, α = 0 for the mirrored surfaces in Fig 1, and pd = 0.4, p_s_ = 0 for the matte images in Fig 2.

The 3D shapes of objects #1-6 were randomly generated with spherical harmonics. To incrementally increase the complexity of the 3D shapes, the maximum degree of the spherical harmonics was limited to 5 for objects #1-2, 7 for objects #3-4, and 10 for objects #5-6. The weights of the spherical harmonics were determined by a random number and normalized so that the power of each degree is inversely proportional to the degree (pink noise). Then the maximum amplitude of the spherical harmonics was normalized to 0.5. The object’s radius of each angle is given by the sum of 1 and the value of the spherical harmonics. The 3D shapes of objects #7-12 were human-made and used in our previous electrophysiological studies of gloss perception [54, 55].

We extracted the orientation field as follows. The image orientation θ(x,y) is the angle that maximizes the magnitude of response p of the oriented filter (first-derivative operator) as *θ(_x_, y)* = argmax *p*^2^ *(θ’(x, y)*). Image anisotropy α(x,y) is defined by the ratio of the minimum and maximum magnitudes of the oriented filter response with respect to its angle [12] as 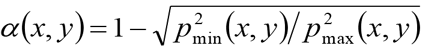 means that the local image is isotropic, and α=1 means that it only consists of one directional component. The steerable pyramid [56, 57] (matlabPyrTools, https://github.com/LabForComputationalVision/matlabPyrTools) was used to extract the image orientation in accordance with previous studies [12, 13, 24]. Responses were obtained by steering the filter through 120 equal orientation steps between 0 and 180°. The finest orientation responses were extracted in accordance with a previous study [12]. Then the amplitudes, which are the squared responses, were downsampled to 256 × 256 pixels and convolved by a 3 × 3 constant filter for noise reduction. Then the image orientation and the image anisotropy were obtained based on the above equations.

We obtained the vertical polarity of intensity gradient pv(x,y) by extracting the sign of the oriented filter response of the vertical direction (θ = 0°) as *Pv* (*X,Y*) = sgn *(p_θ=_0 {x, y)*). The steerable pyramid was used to extract the vertical polarity. The responses of the pyramid level of 256 × 256 resolution were extracted (a relatively low-frequency component compared to the original image resolution). The response values near the boundary are unreliable because they are affected by the image outside of the object region. Therefore, we overwrote the response values within five pixels from the boundary to zeros and smoothed them by a Gaussian filter whose standard deviation is four pixels to reduce the noise and the high-frequency components of the specular reflection.

We derived the signs of the apparent curvature of the image contour as follows. First, we drew a circle centered at a boundary point with a radius of 128 pixels (1/8 of the image size); second, we determined that the curvature sign value at that boundary point is +1 or −1 when the object region’s area within the circle is smaller or larger than the area of the outside object region within the circle; third, for noise reduction, we smoothed the curvature sign values by convolving a constant circular filter of a radius of 16 pixels (1/64 of the image size) and downsampled it to 256 × 256 pixels; then we extracted the signs. The resultant curvature signs of the image contour are shown in S18 Fig.

### Curvature formulation

We described the surface shape of objects by Hessian matrix H(z) of surface depth z(x,y). Because the Hessian matrix is symmetric, H(z) is diagonalized with rotation matrix R as

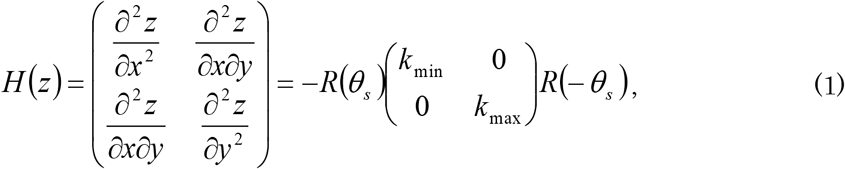

where kmax and kmm are the eigenvalues of the larger and smaller magnitudes. θs, which indicates the angle of the small surface second derivative, is called the surface orientation. There is a minus at the beginning of the right-hand side of Eq 1 so that the surface second derivatives become positive in the case of convex shapes (e.g., sphere). In this study, we described the surface curvature by Hessian matrix based on the image coordinate system instead of the standard curvature that is defined on the object surface’s intrinsic coordinate system. This difference was previously scrutinized [24]. The reason why we adopted the former is that orientation field depends on the Hessian matrix, not on the standard curvatures. For example, in the case of a sphere, the standard curvature is the same at every point on its surface. In contrast, the second derivatives are large near the boundary and small at the center, and correspondingly, the image orientation of the specular reflectance is clear near the boundary and not clear at the center (see Fig 16 of [24]).

Next we introduce other variables and transform the equation. First, surface anisotropy α_s_ is defined as 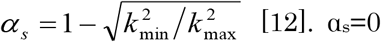 denotes that the magnitude of the two surface second derivatives is the same (e.g., a convex sphere, a concave sphere, or a saddle), and α_s_=1 means that the small surface second derivative is zero (e.g., a convex cylinder or a concave cylinder). Second, variables are introduced so that the surface second derivative’s sign and magnitude are separately described. The sign of the large surface second derivative is represented as σmax e {+1, −1}. +1 and −1 correspond to convex and concave shapes. The magnitude of the large surface second derivative is represented as ka=|kmax |. The sign of the small surface second derivative is represented as σmin e {+1, −1}. Using these variables, the surface second derivatives are described:

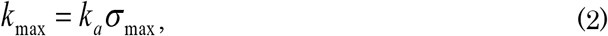

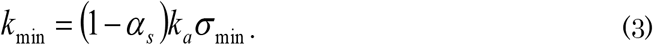

### Relationship between vertical polarity and surface second derivative signs

With the prior knowledge that the object is illuminated from above, we can derive the relationship among the vertical polarity, pv, and the surface second derivative signs. In the case of the Lambert reflectance, the surface luminance is proportional to the inner product of the lighting direction and the surface’s normal direction. Here we assume that the illumination map is stronger as it gets closer to just above (x,y,z)=(0,1,0). As a result, the surface luminance becomes stronger as the surface slant (-dz/dy) is increased. By taking a derivative of this relationship with respect to y and taking the sign, the following equation is obtained:

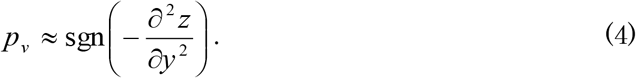

Here we described it as nearly equal instead of equal because the two assumptions of the Lambert reflectance and the just above illumination do not strictly hold in real situations. For example, for images taken outdoors, the angle of the sun (dominant illumination) changes based on time.

We transform Eq 4 into a more available form. The following equation is derived from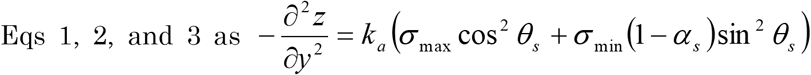Then we used the approximation of orientation θ ~ θs and anisotropy α ~ αs:

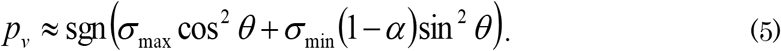

We divided object region Ω into two regions: COS *θ* > (l — $)sin *θ* holds in Ωa, but not in Ωb. Then the following relationship is obtained:

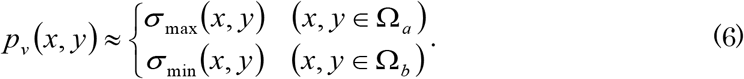

The approximation of Eq 6 was evaluated in our experiment and summarized in the right half of Table 1. All of the results of the objects in the glossy and mirror conditions are shown in S4-S5 and S8-S9 Figs.

### Formulation of cost function

Cost function E consists of two terms: the second derivative constraint given by orientation field C and boundary condition B:

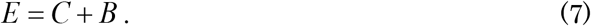

We first explain second derivative constraint C and then boundary condition B, which consists of the following three terms:B=B_0_+B_1_țB_C_.

The second derivative constraint is based on the relationship between the orientation field and the surface second derivatives where the image orientation approximates surface orientation θ ~ θs and the image anisotropy approximates surface anisotropy α ≈ α_s_ [12, 24]. These relationships are described with error terms as *θ_s_ = θ + δθ* and *&_s_ = &* + *δ(X*. These errors were evaluated in our experiment and summarized in then left half of Table 1. For more information, a previous study [24] assessed the orientation error, which depends on the surface anisotropy and the difference between the surface orientation and the illumination map’s orientation. By substituting these equations into Eq 1, we obtain

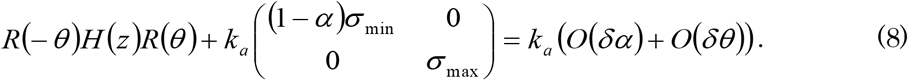

To simplify this equation, we introduce the coordinate axes (u, v) by rotating the original axes (x, y) by image orientation θ(x,y). Note that the axes (u, v) depend on each position based on the image orientation in that position. Then this equation is described as

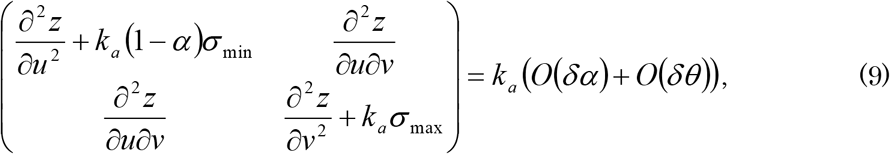

which indicates that the surface strongly bends toward the v direction (the orthogonal direction of the image orientation) by second derivative magnitude ka with sign σmax and the surface weakly bends toward the u direction by second derivative magnitude ka(1 -α) with sign σmin. Second derivative constraint C is based on Eq 9 where the left-hand side is small. The cost is the sum of the squared Frobenius norm of the left-hand side of Eq 9 throughout the object region:

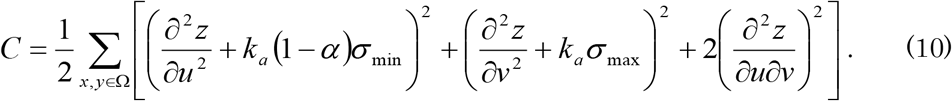

Since this cost function is a quadratic function with respect to z and ka or with respect to σmax and σmin, it is relatively easy to optimize.

Here, because the right-hand side of Eq 9 is proportional to ka, it would be more appropriate to use a cost function that is the sum of the amplitude of the left-hand side of Eq 9 after multiplied by 1/ka. We denote this cost function as C’:

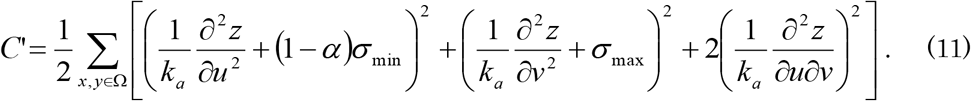

However, cost function C’ is more difficult to optimize. Therefore, we use the first cost function C to obtain the solution, and then with the solution as an initial value, we obtain the improved solution with the second cost function C’. The summarized formula and the minimization of the second cost function are described in Supplementary Note 4 in S1 Text.

Boundary conditions B0 and B1 were introduced to resolve the solution’s ambiguity. B0 resolves the translation ambiguity along the z axis by making the mean depth value zero at the boundary region:

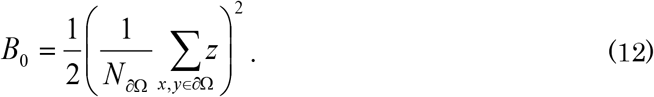

Another ambiguity exists about affine transformations [32]. B1 is introduced so that the solution is not slanted in both the x and y directions:

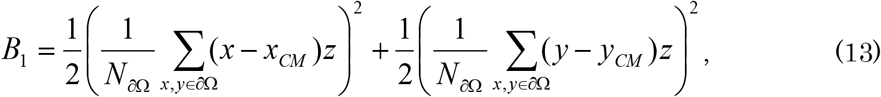

where *^x^cm* and *^y^CM* are the average values of x and y in boundary region dΩ. We summarize these boundary conditions as 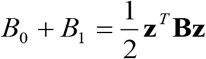 where **z** is the column vector of size Nω × 1 that consists of z(x,y) in object region Ω and B is the coefficient matrix of size NΩ × NΩ.

Next the constraint from the contour was introduced. Assuming that the 3D surface near the boundary is smooth and differentiable, the second derivative toward the normal direction of the contour at the boundary is minus infinity. Therefore, the surface orientation is parallel to the contour and σmax = +1. Moreover, a previous study [58] proved that the sign of the 3D curvature parallel to the contour (=σmin) equals the sign of the apparent curvature of the 2D contour. The apparent curvature sign of the image contour, which is calculated and utilized as the initial values of σmin near the boundary, is also incorporated in the cost function:

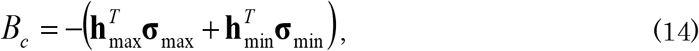

where hmm is a column vector that consists of the contour’s curvature sign (S18 Fig), hmax is a column vector that consists of +1 (near the boundary, where the value exists in S18 Fig) and 0 (otherwise) and σmax and σmin are column vectors that consist of σmax(x,y) and σmin(x,y).

The cost function is summarized as

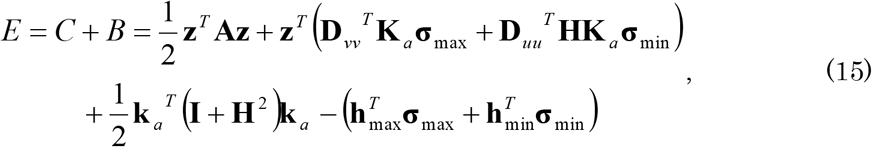

where ka and α are column vectors that consist of ka(x,y) and α(x,y); Ka and H are diagonal matrices with diagonal elements ka and (1-α); D is a matrix that represents the second order differential operator with respect to subscript variables;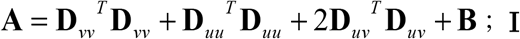 **I** is an identity matrix of size NΩ × N©.Optimal 3D shape z minimizes the cost function. Therefore, the derivative of the cost function with respect to z should be zero. The solution is obtained as

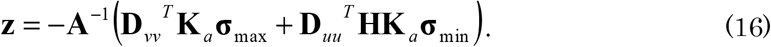

Here, matrix A is invertible since A is positive definite, which can be easily shown. First, the eigenvalue of A is non-negative from the definition (Eqs 7, 10, 12, and 13). Second, there is no zero eigenvalue because of the boundary condition (Eqs 12 and 13). By substituting the solution Eq 16 into Eq 15, the cost function becomes a function of σmax, Omin, and ka:

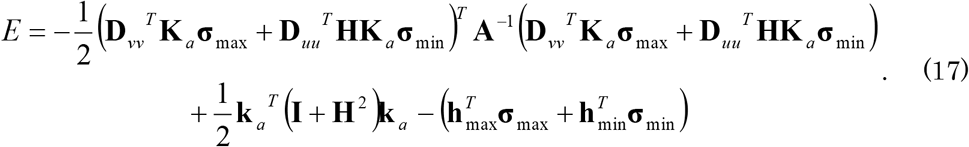

The procedure for minimizing the cost function is described in Supplementary Note 3 in S1 Text.

### Evaluation of recovered depths

We quantify the shape recovery performance by taking the correlation between the recovered depth and the true depth. Note that here we apply the affine transformation so that the slant of the true surface depth becomes zero before taking the depth correlations. The proposed algorithm generates a shape whose slant is zero because of the boundary condition (Eq 13). Therefore, we compared the recovered shape with the true depth after the affine transformation. We summarized the depth correlations without the affine transformation in Supplementary Note 5 in S1 Text.

We used two depth correlations: global and local interior. The global depth correlation is simply the correlation coefficient of the recovered and true depths throughout the object region. However, the global depth correlation tends to become high as long as the depth around the boundary is small, because the true depth is generally very small around the boundary and modest inside the object region. In other words, it is sensitive to the depth around the boundary and insensitive to the details of the shapes inside the object region. Therefore, we proposed a local interior depth correlation, which was calculated as follows. First, we drew a grid that divided the vertical and horizontal axes of the image region into eight (at 32 pixel intervals). Second, we drew a circle centered at an intersection of the grid with a radius of 32 pixels. Third, we measured a depth correlation in the intersection of the circle and the object area after removing the area near the boundary (within 24 pixels from the boundary). We did not measure a depth correlation if the intersection area was smaller than half of the circle’s area. Fourth, we averaged the depth correlation values. As a result, the local interior depth correlation is not affected by the shapes near the boundary and is sensitive to the agreement of the concavity and the convexity inside the object region. Note that we did not evaluate the local interior depth correlation for objects #9 and #11. No depth correlation values were obtained with the above procedure because most of the object region is near the boundary, and the global depth correlation seems sufficient as a measure because there is no fine shape structure inside these object regions.

### Psychophysical experiment

Five unpaid volunteers participated in the experiment (three males and two females; age range, 33-58), all of whom had normal or corrected-to-normal vision and were naïve to its purpose. The experiment was approved by the Ethics Committee for Human Research of National Institute for Physiological Sciences. The experiment was conducted in accordance with the principles of the Helsinki Declaration. Written informed consent was obtained from all participants.

Stimuli were presented on a 58.1 × 38.6 cm flat screen OLED monitor at a distance of 60 cm in a darkened room. Each image subtended at about a 10° visual angle. The stimulus images are shown in Fig 8, although the red crosses in it were not displayed during the experiment. The images of Fig 8A and 8B were rendered by Radiance software with the surface reflection property pd = 0.1, ρ_s_ = 0.15, α = 0 under illumination environments of the Devebec dataset (Galileo’s Tomb for Fig 8A and Eucalyptus Grove for Fig 8B)

Subjects performed two tasks. Both were two-alternative forced choice tasks with no time limits. First, we presented either the image of Fig 8A (Galileo illumination condition) or Fig 8B (Eucalyptus illumination condition). Unfilled, 2.7-cm diameter gray circle centered at the red cross position was superimposed in the first task. Subjects were asked whether the local surface indicated by the circle was convex or concave. Next, we presented the same image and the recovered depth map by the proposed algorithm and the true depth map. The image was located in the center, and the two depth maps were located at the image’s left and right. The left and right arrangements of the recovered and the true depth maps were random. Subjects were asked whether the recovered 3D shape or the true 3D shape more closely resembled the perceived 3D shape from the image. They sequentially performed two tasks for two conditions: the Galileo illumination condition and the Eucalyptus illumination condition. The order of the conditions was counter-balanced among the subjects (two subjects performed the Galileo illumination condition first and three performed the Eucalyptus illumination condition first). Before the experiment, the subjects performed a practice trial with sphere images rendered under another illumination environment (Campus at Sunset of the Devebec dataset) and were instructed about the depth map’s meaning.

## Acknowledgments

The authors thank Masataka Sawayama, Shin’ya Nishida, Yinqiang Zheng, and Imari Sato for their helpful comments. This research was supported by a contract with the National Institute of Information and Communications Technology entitled, ‘Development of network dynamics modeling methods for human brain data simulation systems’ (grant #173), and Grant-in-Aid for Scientific Research on Innovative Areas “Shitsukan” (15H05916 and 25135737) from MEXT, Japan, and the ImPACT Program of the Council for Science, Technology and Innovation (Cabinet Office, Government of Japan).

